# Synergistic Effects of pH and Temperature on Dengue Virus Envelope Dimers: Insights from Microsecond Molecular Dynamics Simulations

**DOI:** 10.1101/2025.10.04.680450

**Authors:** Georcki Ropón-Palacios, Ingrid B. S. Martins, Walter Rocchia, Alexandre S. de Araujo

## Abstract

Dengue pathogenesis depends on a conformational rearrangement of the envelope (E) glycoprotein induced by low pH; however, how pH and temperature cooperate to destabilize its prefusion dimeric state, a fundamental step that enables the formation of the fusion-active trimer, remains unclear. Here, we investigate this synergy for Dengue serotypes of greatest medical relevance, DENV–2 and DENV–3. All–atom molecular dynamics simulations were performed on the microsecond scale, at 28 °C, 37 °C, and 40 °C under pH 5–7, using CHARMM36m force fields. The increase in temperature from 28 °C to 37 °C doubled the conformational space explored at neutral pH and weakened both dimers through the reduction or loss of the interaction network along the dimeric interface. Acidification amplified this effect in a serotype–specific manner: DENV–2 required pH 5, while DENV–3 responded at pH 6. Distance and principal component analyses revealed an asymmetric dissociation route, termed the “compensatory embrace”, in which retraction of Domains I and III in one monomer is balanced by advancement of its partner on the opposite side, temporarily preserving the number of inter–subunit contacts. Structural analyses highlighted the difference in histidine distribution between the serotypes. These results outline hierarchical physicochemical triggers that convert the dimer into fusion–competent monomers. Targeting the interfacial interaction network or reinforcing the “compensatory embrace” to prevent completion of dissociation offers new perspectives for broad–spectrum antivirals and immunogen design, underscoring the value of long–timescale molecular dynamics in drug discovery.

## 1 INTRODUCTION

Dengue virus (DENV) remains the most prevalent arthropod-borne pathogen worldwide, with 390 million infections per year and 96 million clinically apparent cases according to a landmark modelling study ^1^. Disease outcomes range from self-limited febrile illness to dengue hemorrhagic fever and shock syndrome^2,3^. A major driver of severe manifestations is antibody-dependent enhancement (ADE), whereby sub-neutralizing antibodies from prior exposure or vaccination facilitate Fc–mediated uptake of a heterologous serotype^4–6^. The DENV envelope glycoprotein (E) orchestrates both cell receptor binding and the low–pH membrane–fusion reaction that delivers the viral genome into host cells^7,8^.

Structurally, two E–protein ectodomains (pE) assemble into an antiparallel homodimer that lies flat on the mature virion surface^9–11^. Upon endosomal acidification, the dimer is destabilized, exposing a hydrophobic fusion loop in Domain II that inserts into the target membrane^12–17^. Recent cryo–EM snapshots have captured pH–triggered intermediates, underscoring the ectodomain’s conformational plasticity^10,11,18^. Temperature is an equally critical variable: mosquito-like, physiological, and febrile conditions (28 °C, 37 °C, and 40 °C) modulate virion “breathing,” alter epitope exposure, and weaken E-dimer stability in a serotype-dependent manner, this includes temperature-driven structural transitions captured by cryo-EM and HDX-MS, and a strong shift toward monomeric E at 37 °C that diminishes quaternary-epitope display^18–21^.

Vaccine development is complicated by the existence of four antigenically distinct serotypes (DENV-1–4) and by the need to elicit balanced immunity^22–24^. The quaternary E–dimer epitope (EDE)—which bridges both protomers—has emerged as a prime target for broad neutralization^25,26^. Yet most structural and computational analyses have focused on single serotypes or on acidification in isolation, leaving the combined effects of pH and temperature on intact E–protein dimers largely unexplored.

Here, we close this gap using microsecond all–atom molecular dynamics (MD) simulations, principal component analysis (PCA), and free-energy landscape (FEL) mapping to interrogate the epidemiologically dominant serotypes DENV–2 and DENV–3 across pH 5–7 and 28 °C, 37 °C, 40 °C. Our results reveal that (i) neutral conditions perturbed the structural stability, flexibility and expand the conformational space in DENV–2 e DENV–3 when the temperature is switched from 28 to 37 and 40 °C (ii) an acid- and heat–dependent “compensatory embrace” that transiently preserves the number of inter– subunit contacts, a key process to be targeted, (iii) a temperature– and pH–dependent weakening of the interfacial interaction network, and (iv) serotype–specific histidine networks that could fine–tune the activation threshold. These insights provide a physicochemical roadmap for designing antivirals and immunogens aimed at stabilizing protective dimer states or disrupting pathogenic transitions.

## 2 MATERIALS AND METHODS

### System Preparation

The atomic coordinates of dengue virus E protein dimers (serotypes DENV–2 and DENV–3; PDB IDs 1OAN and 7A3S, respectively) were retrieved from the Protein Data Bank ^27–29^. Histidine protonation states at pH 5, 6, and 7 were assigned using PROPKA3^30^ through the PDBReader module of CHARMM-GUI^31–33^, which also reconstructed missing loops and unresolved atoms. All-atom topologies (PSF) and coordinates (PDB) were generated with the CHARMM36m force field including CMAP corrections ^34,35^. Each dimer was solvated in a TIP3P water box extending 15 Å beyond the solvent–accessible surface. To reproduce physiological conditions, Na^+^ and Cl^−^ ions were added to neutralize the systems and achieve an ionic strength of 150 mM using the MSTBx v0.5 toolkit (available at: https://github.com/SSiLiq/MSTBx). The final systems comprised approximately 600 thousand atoms for both DENV–2 and DENV–3, including explicit water molecules and counterions, with box dimensions of 184 *×* 184 *×* 184 Å and 182 *×* 182 *×* 182 Å, respectively.

### Molecular Dynamics Simulation

All simulations were performed with NAMD 2.14/3.0b6^36,37^ following a consistent equilibration and production protocol. Each system was first subjected to 2,000 steps of energy minimization using the conjugate gradient algorithm, with positional restraints of 5 kcal mol^−1^ Å^−2^ applied to backbone atoms to relieve steric clashes. Equilibration was carried out in two NPT stages: (1) 2 ns of gradual heating from 60 K to the target temperature (301, 310, or 313 K; approximately 28, 37, and 40 °C) in 1 K increments, while maintaining pressure at 1.01325 bar with a Langevin thermostat and a Nosé–Hoover Langevin piston barostat; (2) 3 ns of NPT at the final temperature and 1.01325 bar under the same restraints and thermostat–barostat settings. Production dynamics were then conducted for 1 µs per system in the NPT ensemble. During equilibration and production, a 2 fs integration timestep was employed. Long–range electrostatics were evaluated every two steps (fullElectFrequency = 2), and nonbonded interactions every step (nonbondedFreq = 1). All covalent bonds involving hydrogens were constrained (rigidBonds = all). Long–range electrostatics were computed using Particle Mesh Ewald with a 1 Å grid spacing^38,39^. Short–range van der Waals interactions were truncated at 12 Å with force switching enabled and a switching onset at 10 Å. The Langevin damping coefficient was set to 1 ps^−1^, and piston oscillation and decay periods were set to 200 fs and 100 fs, respectively. Trajectories were saved every 50 ps (20,000 frames per µs) for all conditions: DENV–2/DENV–3, pH 5/6/7, and temperatures of 301 K (28 °C), 310 K (37 °C), and 313 K (40 °C).

### Trajectory Analysis

Trajectory processing was performed with VMD 1.9.4a57 and MDAnalysis 2.7^40–42^, supplemented with in–house Python, Tcl, and Bash scripts (available at: https://github.com/groponp/CompBiology-Biophysics). Structural stability was assessed by calculating the root–mean–square deviation (RMSD) of *C*_*α*_ atoms relative to the initial configuration. Interface RMSD was computed using only *C*_*α*_ atoms within 8 Å of the opposite monomer, and per-residue flexibility was quantified using the root–mean– square fluctuation (RMSF). Dimer dissociation events were defined as frames in which the center–of– mass distance between the monomers’ *C*_*α*_ atoms exceeded 60 Å. Conformational dynamics were further explored by principal component analysis (PCA). *C*_*α*_ trajectories were aligned to the reference structure, and the covariance matrix was constructed to extract the first two principal components (*PC*1 and *PC*2), which together captured the largest fraction of atomic variance. Free energy landscapes (FELs) were then computed as:

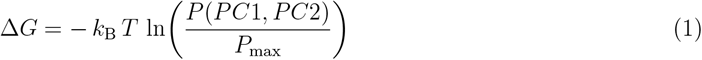

where *k*_*B*_ is the Boltzmann constant, *T* the absolute temperature, *P* (*PC*1, *PC*2), the joint probability distribution of *PC*1 and *PC*2, and *P*_max_ the maximum probability value (i.e., the reference state). For comparison purposes, the FEL minimum of each system was shifted to zero, corresponding to the global lowest free energy state.

Interface contacts were defined as residue pairs with *C*_*α*_ atoms located within 8 Å across the dimer interface. Contact frequencies were computed for the simulations at 28, 37, and 40 °C at each pH, and only contacts present in all three temperatures for a given pH were retained. GetContacts (available at: https://github.com/getcontacts/getcontacts) was then used to classify interactions into specific types (e.g., hydrogen bonds, *π*–cation interactions, and salt bridges).

The figures were rendered using VMD 1.9.4a57, ChimeraX 1.8, and Matplotlib 3.7.1^40,43–45^.

## 3 RESULTS AND DISCUSSION

In this section, we present and discuss our findings on how pH and temperature jointly modulate the structural dynamics of the dengue virus ectodomain envelope protein (pE) dimer in serotypes DENV–2 and DENV–3. Our data show that acidic pH and elevated temperature remodel the conformational landscape of the E protein, indicating potential alterations in its ability to mediate host cell entry and evade the host immune system, while also revealing possible targets for the action of small–molecule inhibitors of its fusogenic activity. Elucidating these physicochemical triggers is therefore critical for the rational design of potent neutralizing antibodies, antiviral drugs, and next–generation vaccine candidates.

### Structural Comparison of the pE Ectodomain and Histidine Distribution in DENV–2/3

In Figure 1A, we show the high-resolution X-ray crystal structures of the E protein ectodomain for DENV–2 (PDB ID 1OAN) and DENV–3 (PDB ID 7A3S) side by side in cartoon representation. Domains I (red), II (yellow), III (blue), and the fusion loop (purple) were highlighted, and all histidines were rendered as sticks and labeled by residue number.

**Figure 1.**
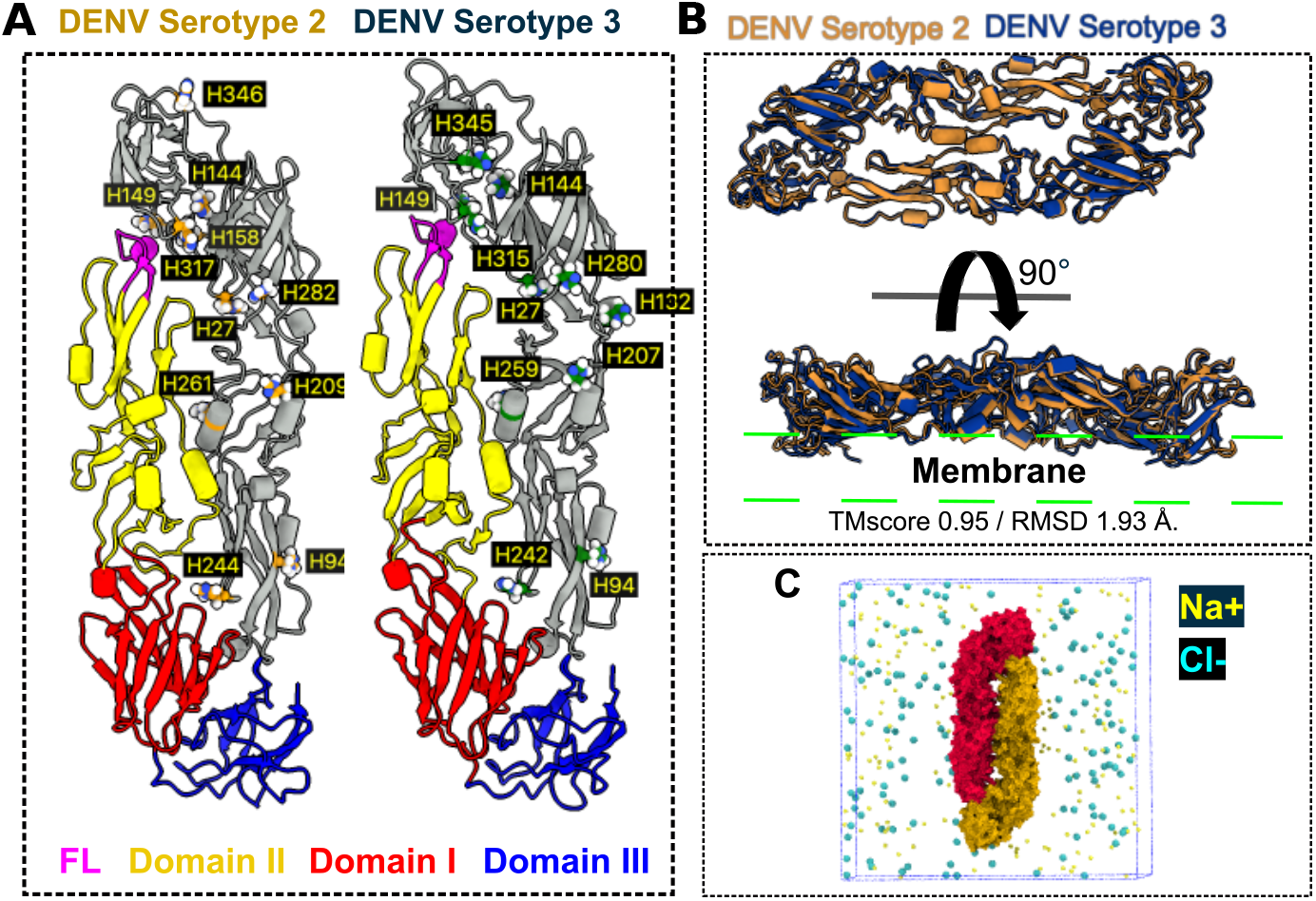
Experimental models of the E protein (pE) from DENV–2 and DENV–3 serotypes. (A) Complete crystallographic structures of the ectodomain, with the DENV–2 and DENV–3 dimers displayed side by side. Domains I (red), II (yellow), and III (blue), as well as the fusion loop (purple), are highlighted in distinct colors. All histidines are shown as sticks and labeled with their corresponding residue numbers. (B) Structural superposition of the ectodomains of DENV-2 and DENV-3. TM–score and RMSD values were obtained with the TM-align program ^46^. (C) Solvated model of pE. Na^+^ and Cl ions are shown as van der Waals spheres; water molecules were omitted for clarity.

Our analysis reveals subtle differences in the spatial arrangement of histidines along the protomer, in agreement with previous reports^47^. Both serotypes conserve four histidines adjacent to the fusion loop: His144, His149, His158, and His317 in DENV–2, and His144, His149, His315, and His345 in DENV–3; among them, His144 and His317 (His315 in DENV–3) have been highlighted in the literature as major pH sensors at the DI–DIII interface^13,47^. Although His149/His158 in DENV–2 and His149/His345 in DENV–3 lack direct experimental validation as essential pH sensors, we hypothesize that these secondary histidines, together with the others, may fine–tune the acid–dependent activation threshold for dimer dissociation in a serotype–specific manner without perturbing the global fold. This effect likely arises through a network of interactions around the fusion loop, as suggested in the literature^48^. For example, His244 in DENV–2 (His242 in DENV–3) has been proposed to mediate interactions with Asp63 of prM and to play a role in viral maturation rather than fusion^49,50^.

We also superimposed the two ectodomains using TM-align (Figure 1B), yielding a TM–score of 0.95 and an RMSD of 1.93 Å, which confirms strong conservation of the overall fold despite 25–40 % sequence divergence between serotypes ^51–53^. A closer examination of the structures–particularly the surface–exposed loops—revealed that most of the structural variability is localized to Domains I and III, whereas Domain II remains largely unchanged^54^. We propose that these subtle structural differences underlie reported variations in thermal stability, immunogenicity^52,55^, receptor affinity, and potentially dimer stability and dynamics^52,56^. Thus, even with a globally conserved fold, minor primary– sequence divergences pose challenges for the rational design of vaccines, neutralizing antibodies, and small-molecule inhibitors.

Finally, Figure 1C illustrates the solvated crystallographic systems we prepared for our MD simulations. Each protomer is shown in a distinct surface color; Na^+^ and Cl^−^ ions appear as van der Waals spheres within the dashed simulation box; and water molecules has been omitted for clarity. This structural framework underpins our dynamic analyses, enabling us to correlate specific conformational responses with variations in pH and temperature.

### Dimer Stability under pH and Temperature Variations

To assess how pH and temperature jointly influence the stability of the pE dimer in DENV–2 and DENV–3, we performed all–atom molecular dynamics simulations at 28 °C, 37 °C and 40 °C across pH 5, 6 and 7. These serotypes were chosen for their epidemiological relevance ^57–59^ and for subtle primary-sequence variations that, as confirmed in Figure 1B, do not alter the global ectodomain fold but may drive distinct dynamic behaviors. In Figure 2A, each monomer is rendered in a distinct color and both frontal and posterior views of the dimer are shown.

**Figure 2.**
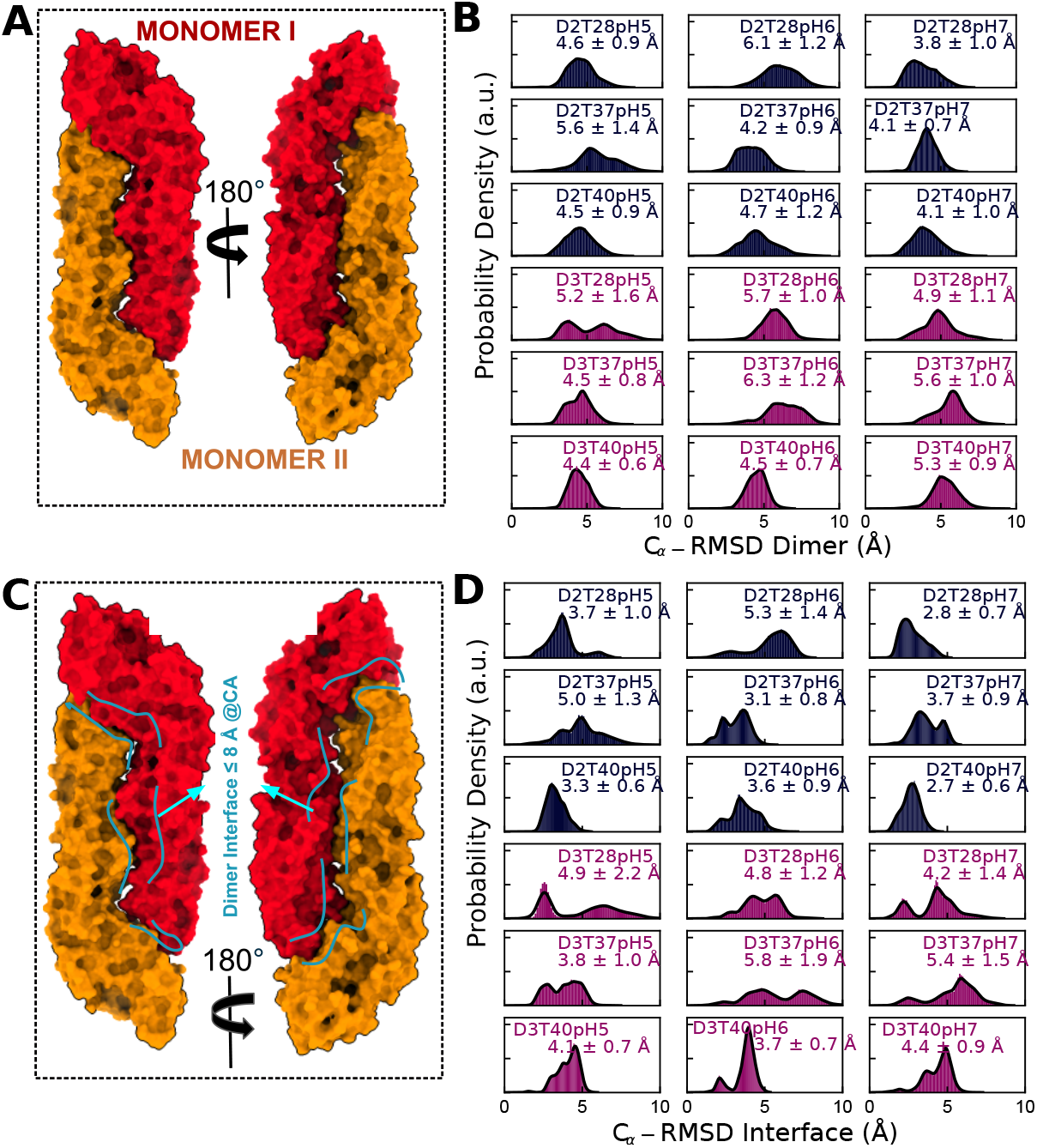
Conformational stability of the E protein from DENV–2 and DENV–3 serotypes. (A) Graphic representation of the pE dimer, with each monomer highlighted in a different color and shown in both front and back views. (B) Global *C*_*α*_ RMSD histograms, calculated from simulations performed at 28 °C, 37 °C, and 40 °C under pH 5, 6, and 7. (C) Interface selection diagram indicating the residues whose *C*_*α*_ atoms are within *≤* 8 Å of the other protomer. (D) Interface RMSD histograms (*C*_*α*_) for the residues considered in panel (C), under the same temperature and pH conditions.

Our global RMSD histograms (Figure 2B) reveal the combined influence of pH and temperature on dimer stability. At pH 5, DENV–2 reaches its highest RMSD (5.6 ± 1.4 Å) at 37 °C—an endosomal temperature typical of the human host^50^—consistent with fusion of the NGC DENV–2 strain peaking at pH 6.2 or below^50,60^. In contrast, DENV–3 exhibits its maximum RMSD (6.3 ± 1.2 Å) at 37 °C only at pH 6, aligned with experimental reports ^61^ and underscoring how differences in histidine distribution tune each serotype’s pH sensitivity. Still at pH 6 we observe for DENV–2 peaks at 28 °C (RMSD 6.1 ± 1.2 Å), reflecting mosquito-like conditions, whereas DENV–3 remains maximal at 37 °C, in agreement with binding studies reporting optimal attachment between 37 °C and 40 °C, with 37 °C being most favorable^62,63^. At neutral pH, DENV–2 maintains elevated RMSD (4.1 ± 0.7 Å) at both 37 °C and 40 °C, while DENV–3’s maximum RMSD (5.6 ± 1.0 Å) occurs only at 37 °C. This behavior concurs with literature indicating that neutral pH and physiological temperatures favor dimer–to–monomer dissociation; under these conditions, DENV–3 displays a greater propensity for the monomeric form^61^.

### Local Interface Fluctuations and Residue Flexibility

To gain deeper insight, we next analyzed the dimer interface, defined as pairs of *C*_*α*_ atoms within 8 Å across monomers (Figure 2C), and extracted interface RMSD histograms (Figure 2D) to capture local contact loosening. For DENV–2 at pH 5 and 37 °C, the mean interface RMSD is 5.0 ± 1.3 Å, confirming significant loosening; at pH 6 and 28 °C, the peak shifts similarly (5.3 ± 1.4 Å), suggesting that early–stage dissociation could occur under mosquito–like conditions ^64^. By contrast, at physiological pH (6–7) and 37–40 °C, interface RMSD remains near 3 Å, but rises to approximately 3.7 Å at pH 7 and 37 °C before settling at 2.7 ± 0.6 Å at 40 °C, highlighting a thermal “switch” between vector and host environments.

In DENV–3, the critical interface RMSD (5.8 ± 1.9 Å) occurs at pH 6 and 37 °C; at pH 5 and 28 °C it remains elevated (4.9 ± 2.2 Å), shifting to 4.4 ± 2.2 Å at pH 7 and 40 °C—indicating that neutral and febrile conditions still preferentially destabilize this serotype. Under physiological pH, raising the temperature from 28 °C to 37 °C reproduces the same thermal “switch” pattern, with RMSD of 4.2 ± 1.4 Å at 28 °C and 5.4 ± 1.5 Å at 37 °C. However, cryo-EM data of mature virions show that soluble DENV-3 ectodomains incubated at 37 °C remain structurally similar to those at 28 °C^65^, suggesting reversible thermal “breathing” response in solution without irreversible unfolding, in contrast to the NGC strain of DENV–2. In light of this apparent paradox, we hypothesize that the thermal “switch” operates differently in soluble dimers versus the virion surface–potentially modulated by viral components such as a protein M and the lipid bilayer and varying by strain–and therefore must be contextualized to the specific structural level under investigation to avoid confusion with the “breathing” in virion context.

In light of the thermal “switch” described above, Figure 3 presents per–residue RMSF profiles for each monomer at 28 °C, 37 °C and 40 °C across pH 5, 6 and 7. While global RMSF values are comparable between serotypes, distinct peaks emerge: at pH 7 and 37 °C, DENV-2 shows elevated flexibility in Domains I and II (residues approximately 100–200), which facilitate fusion-related rearrangements, and at 40 °C in Domains I–III (residues approximately 380–394). Under pH 5 and 37 °C, DENV–2 exhibits its highest RMSF (3.0 ± 1.2 Å) in chain B. In contrast, DENV–3 reaches maximum flexibility (2.8 ± 1.0 Å) in chain B at pH 5 and 28 °C, followed by pH 6 at 37 °C (2.5 ± 1.0 Å) in chain B and pH 7 at 37 °C (2.5 ± 1.0 Å) in chain C, with Domain III consistently the most mobile region. Crucially, this pattern of specific peaks also uncovers an asymmetric behavior: as one monomer’s flexibility increases, the partner’s decreases, indicative of an asymmetric dissociation mechanism that we explore in the next section.

**Figure 3.**
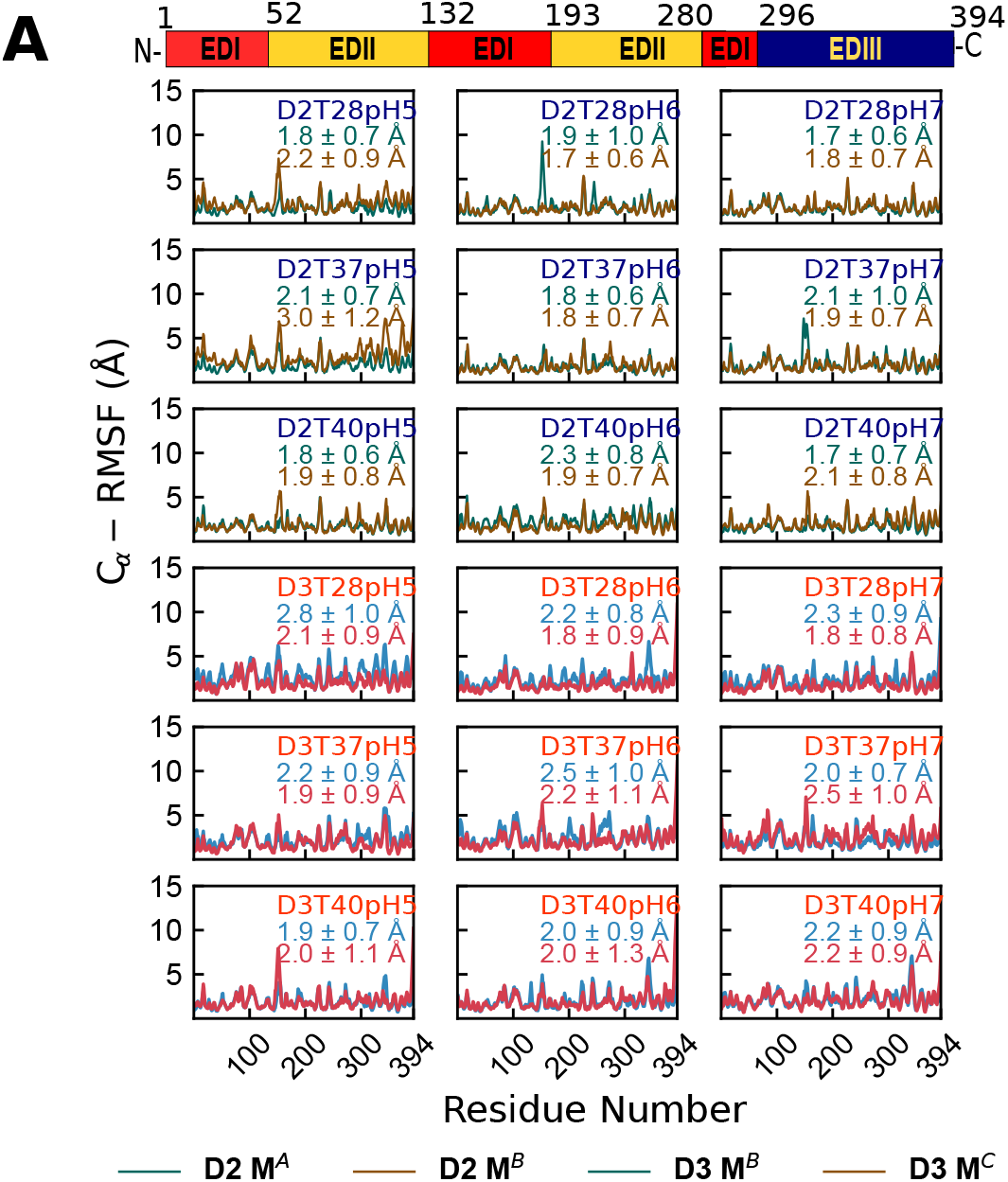
Residual flexibility profile (RMSF) of the E protein from DENV–2 and DENV–3 serotypes. (A) RMSF curves along the sequence for each monomer, obtained from simulations at 28 °C, 37 °C, and 40 °C under pH 5, 6, and 7. Above, a schematic diagram of domains DI–DIII indicates the location of residues to facilitate interpretation. In the legend of the graph, the abbreviations D2 MA, D2 MB, D3 MB, and D3 MC indicate the viral serotype (D2: DENV-2; D3: DENV-3) and the corresponding monomer chain, represented by MX, where X = A, B, C.

### pH and Temperature Trigger pE Dimer Dissociation within 1 µs of Molecular Dynamics

In order to observe the onset of dissociation, we monitored the distance between the centers of mass of the monomers (d_*M*→*M*_) over 1 µs of all–atom molecular dynamics simulations; a duration sufficient to capture key conformational rearrangements ^66^. Based on the crystallographic dimer structure, in which the inter-monomer distance is approximately 50 Å, we defined 60 Å as the threshold for dimer dissociation (Figure 4A, green line). Distances beyond this value indicate the onset of separation, as they exceed the native spacing observed in the crystal. In Figure 4A, we present d_*M*→*M*_ histograms for DENV–2 and DENV–3 under each pH–temperature combination; in Figure 4B, we display representative snapshots of the three trajectories that exceeded this threshold.

**Figure 4.**
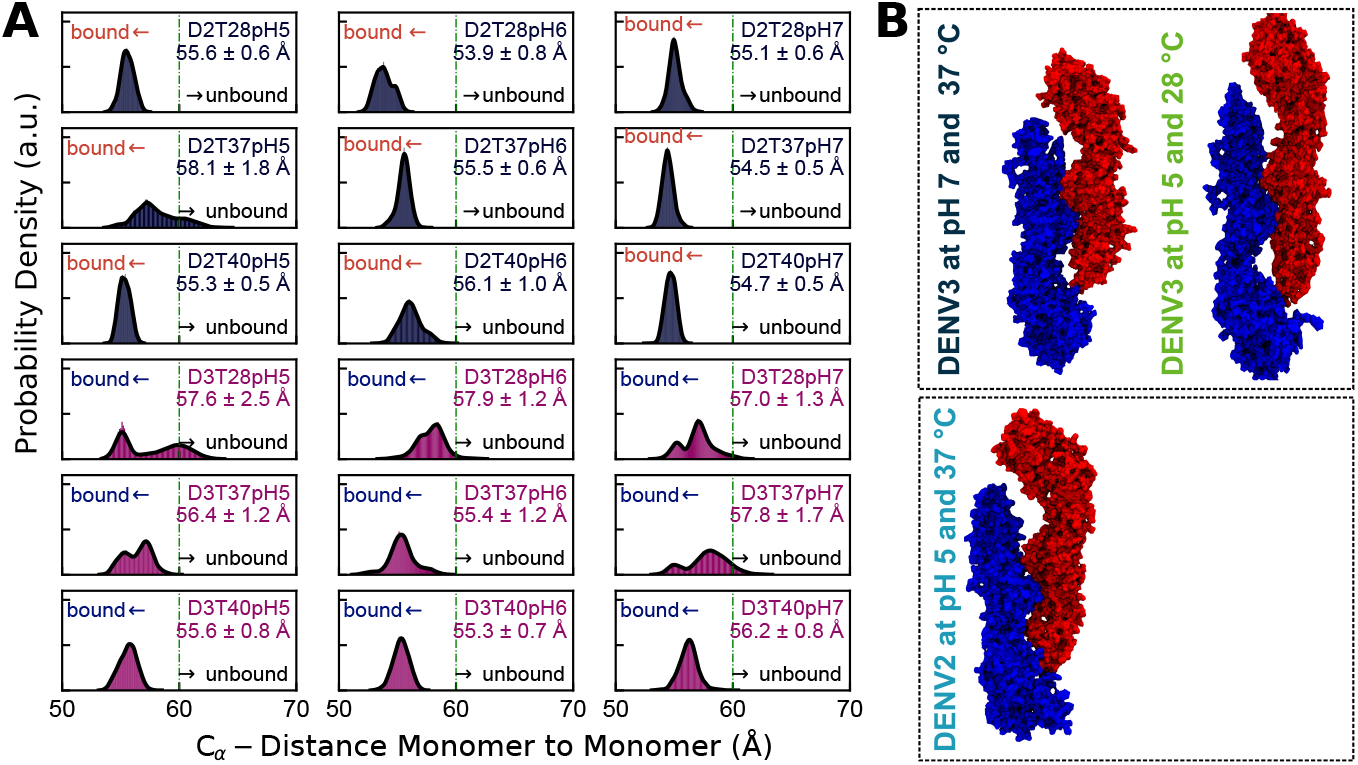
Dissociation dynamics of E protein dimers from DENV–2 and DENV–3 under different pH and temperature combinations. (A) Histograms of the distance between monomer centers of mass (d_*M*→*M*_) obtained after 1 µs of molecular dynamics. The green line marks 60 Å, the threshold value defined as the onset of dimer dissociation. (B) Representative structural snapshots of the three trajectories that exhibited dimer dissociation.

Our results agree with evidence that low pH acts as a trigger for E protein fusion, promoting conformational changes in the ectodomain^47^, while physiological temperatures favor the dimer–to–monomer transition in both DENV and ZIKV^57,61^. By analyzing histograms and three–dimensional structures, We identified a mechanism of asymmetric dissociation: while one monomer loses contacts, with its domains I and III moving away from domain II of the opposite monomer, the other side of the dimer tightens as the adjacent monomer approaches. This rearrangement, which we term a “compensatory embrace”, can preserve or even slightly increase the overall number of inter–subunit contacts, thereby masking partial separation; an effect that is more pronounced in DENV–2 ()(Figure 4B, see Video S1-S3 in the Supporting Information).

However, at mildly acidic pH (pH approximately 6), none of our simulations exceeded the 60 Å threshold, suggesting that this range may act as a conformational restraint, attenuating inter–monomer fluctuations at physiological temperature. Nevertheless, it remains unclear whether this mechanism also applies to the virion, where components such as protein M and the lipid bilayer may modulate it, or whether the absence of dissociation merely reflects sampling limitations in our simulations–especially considering that experimental evidence points to pH approximately 6 as an optimal condition for the virus.

Given the interesting nature of the “compensatory embrace” in preserving inter–subunit contacts, it would be a suitable target to design molecules capable of stabilizing it. This would represent an innovative strategy to inhibit E dimer dissociation and, thereby, block the viral fusion process.

### Effect of pH and Temperature on the Essential Dynamics of the pE Dimer

To elucidate how pH and temperature jointly modulate the essential motions of the pE dimer in DENV–2 and DENV–3, we applied an integrated principal component analysis (PCA) and free energy landscape (FEL) approach (Figures 5A–D). In Figure 5A, each point represents a conformation sampled over 1 µs, colored according to its simulation time. At neutral pH, raising the temperature from 28 °C to 37 °C nearly doubled the conformational space explored: in DENV–2, the originally compact cloud split into two connected clusters, whereas DENV–3 dispersed into three distinct regions. Consistently, the FELs in Figure 5B show that DENV–2 shifts from a single basin of approximately 2.3 kBT at 28 °C to two minima separated by approximately 0.8 k_*B*_T at 37 °C, while DENV–3 displays three shallow basins with internal barriers below 0.5 k_*B*_T.

**Figure 5.**
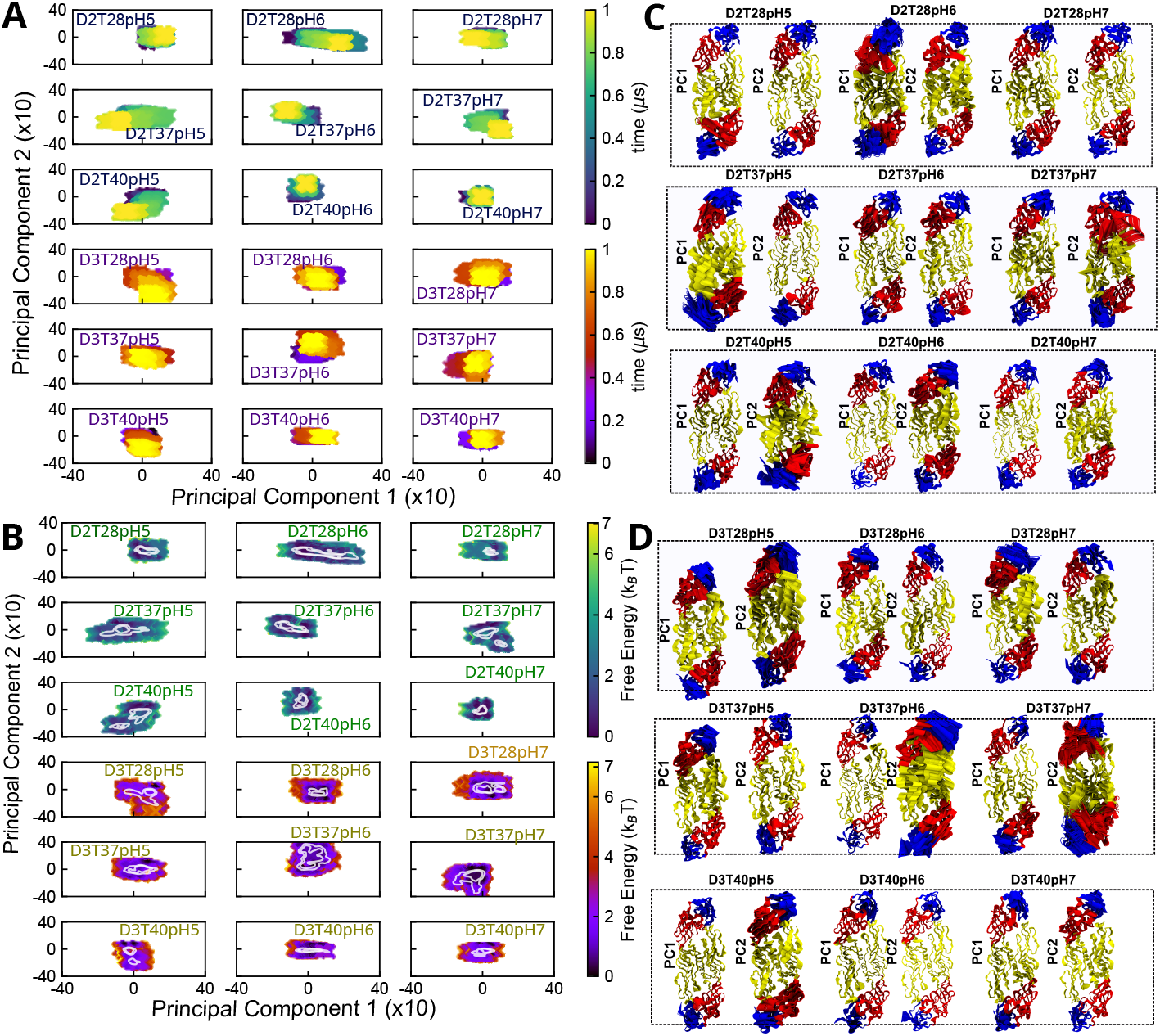
Combined effect of pH and temperature on the essential dynamics of the E–protein dimer from DENV–2 and DENV–3 serotypes. (A) PCA projections; each point represents a conformation sampled over 1 µs, with a color gradient indicating temporal evolution. (B) Free energy landscapes (FEL) on the PC1 *×* PC2 plane. (C) Dominant modes of DENV–2. (D) Dominant modes of DENV–3. In panels C and D, domains DI, DII, and DIII are colored according to the conventional E–protein palette defined previously.

We next examined the extreme PCA vectors (Figures 5C, 5D), which indicate that PC1 is dominated by the rotation of Domain III and PC2 by an asymmetric swing of Domain I around the DI–DIII interface. These motions underpin our “compensatory embrace” hypothesis, in which the retreat of DIII in one monomer is offset by an advance of DI in its partner, thereby preserving inter–subunit contacts even as the dimer approaches monomerization^57^. In DENV–3, this effect is even more pronounced, consistent with the substantial monomeric fraction observed at 28 °C that further increases at 37 °C^61,67^. Such large–scale displacements likely hinder recognition of quaternary epitopes and drive researchers to engineer stabilizing mutations—an insight valuable for antibody design and small-molecule inhibitor development^25,67^.

Strikingly, at neutral pH, raising the temperature from 28 °C to 37 °C amplifies motions in Domains I and II along PC2—an effect more pronounced in DENV–3—which may induce contact loss and favor dimer dissociation during the thermal transition, in line with the high enthalpy (*ΔH*) reported in the literature^61,67^.

At pH 6, we observe a conformational “lock,” which, as noted above, may reflect the absence of additional viral components in the modeled system or limited sampling in our simulations. For DENV–2 at 28 °C, the FEL collapses into a single basin of approximately 1.6 k_*B*_T and the PCA cloud contracts to a radius below 15 Å, with fluctuations confined to Domain I and minimal motion of DIII (Figure 5C).

An identical pattern emerges for DENV-3 at 37 °C, confirming that mildly acidic conditions reduce large–scale rearrangements^68^, yet still perturb dimer stability.

By contrast, acidification to pH 5 at 37 °C remodels the FEL into interconnected shallow minima (*<* 1.2 k_*B*_T) and projects Domain II toward solvent—a prelude to progressive release of DIII (Figures 5A–D). This opening is more pronounced in DENV–2 than in DENV–3. During this process, DI and DII swing in opposite directions to re–establish the “compensatory embrace”; once these motions reach their maximum amplitude, we propose that the interface loses cohesion and the dimer crosses the 60 Å separation threshold (Figure 4), confirming acidic pH as the conformational trigger for dimer dissociation^68^.

### Synergistic Effect of pH and Temperature on Molecular Interactions at the DENV E– Protein Dimer Interface

To elucidate, in detail, how temperature and pH jointly modulate the interaction network at the DENV–2 and DENV–3 E–protein dimer interface, we mapped all interfacial contacts (see Materials and Methods; Figure 6A–B). In Figure 6A, blue arrows highlight those residues whose contact frequency decreases when the temperature is raised from 28 °C to 37 °C, then partially recovers at 40 °C; and a direct increase in temperature from 28 °C to 40 °C also diminishes these contacts. We then identified interactions that occur exclusively at acidic pH (5 and 6) and are absent at pH 7, all measured at 37 °C.

**Figure 6.**
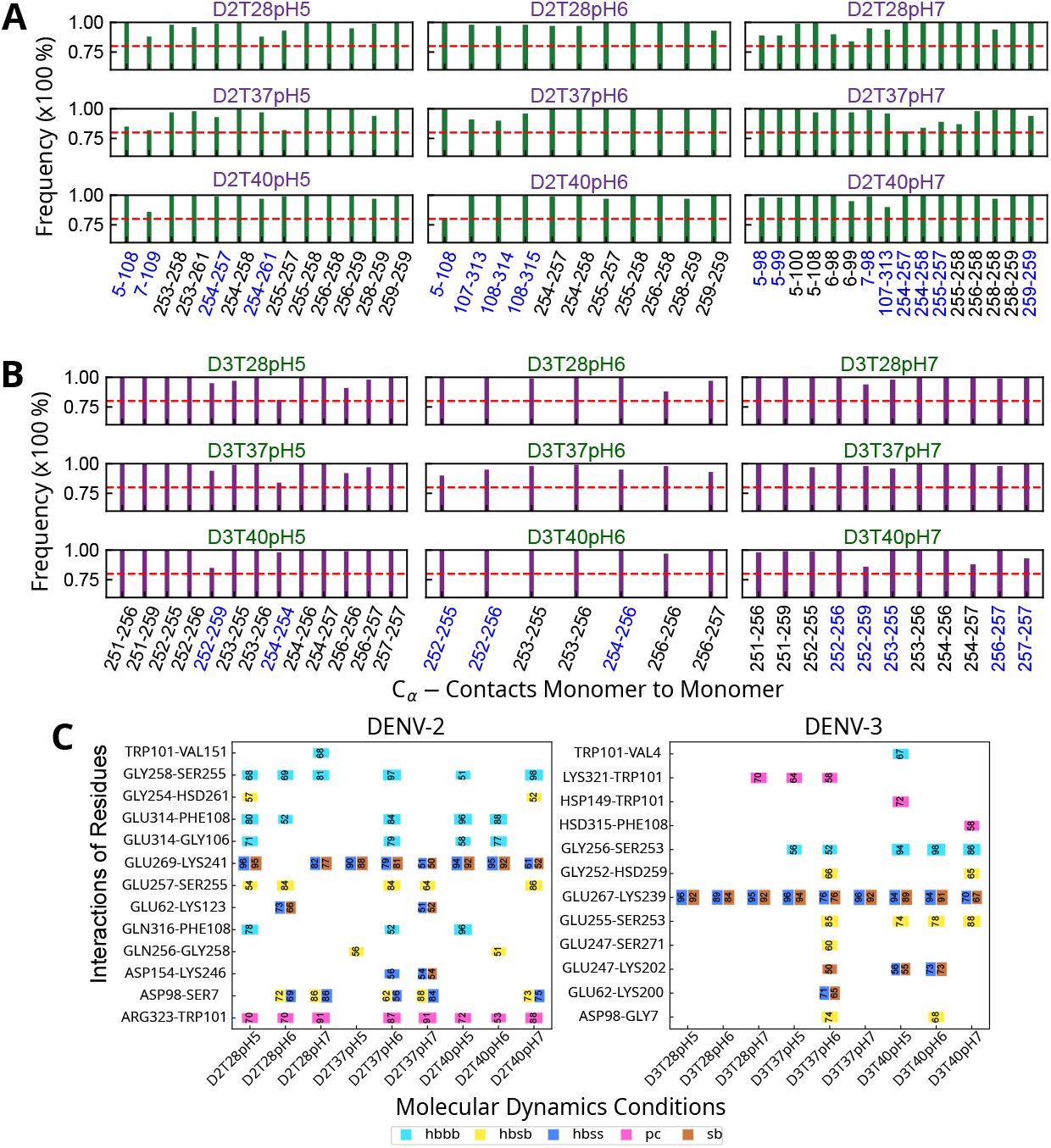
Combined effect of pH and temperature on the molecular interactions of the E–protein dimer from DENV–2 and DENV–3 serotypes. (A) and (B) Frequency of common contacts across different temperatures at a given pH. Blue–colored pairs indicate residues whose contact frequency is sensitive to temperature changes. (C) Quantification in DENV–2 and DENV–3 of backbone–backbone (hbbb), sidechain–backbone (hbsb), and sidechain–sidechain (hbss) hydrogen bonds, as well as *π*–cation (pc) and salt bridges (sb), under different simulation conditions.

Under these conditions, in the DENV–2 dimer the most prominent van der Waals contacts were Gly5–Phe108 and Ser7–Gly109 (DI–DII interface), Gly254–Glu257 (both in DII), and Phe108–Glu314 and Phe108–Thr315 (DII–DIII interface). The latter pair has been targeted for disulfide–bridge engineering to improve dimer stability^69^. In the DENV–3 dimer, the key contacts were Gly252–His259, Gln254–Gln254, Gly256–Gly256, Gly252–Gly256, and Gln254–Gly256 (all within DII). These van der Waals interactions proved potentially critical for dimer stability in solution and, consequently, for virion integrity^61^.

Next, to characterize the underlying molecular mechanisms in these dengue variants, we evaluated and quantified hydrogen bonds–backbone–backbone (hbbb), sidechain–backbone (hbsb), and sidechain– sidechain (hbss)—as well as *π*-cation (pc) interactions and salt bridges (sb). These results are shown in Figure 6C. First, we analyzed the effect of temperature at pH 7 for DENV–2: raising the temperature from 28 °C to 37 °C reduced the Asp98–Ser7 (DII–DI interface) hbss frequency from 86 % to 84 %, falling to below 76 % at 40 °C for both hbsb and hbss. We also found that Asp154–Lys246 (DI–DII) lost its salt bridge and hbss entirely outside the 37 °C condition, retaining only 54 % at that temperature– consistent with the exothermic nature of salt bridges, whose stability decreases as temperature rises ^61^. In the Glu269–Lys241 (DII–DII) pair, we recorded decreases of over 20 % in hbss and sb at 37 °C and approximately 10 % at 40 °C, with the greatest reduction occurring between 28 °C and 37 °C; this pair lies in the intradimer interface (residues 238–260), which shows temperature–dependent disruption in hydrogen–deuterium exchange assays^70^. Additionally, the Gly258–Ser255 (DII–DII) hbbb contact was present only at 28 °C and 40 °C, suggesting a high–temperature conformational rearrangement, while Trp101–Val151 (DI–DI) remained absent at 37 °C and 40 °C, indicating loss of hbbb under those conditions. In the DENV–3 dimer, Glu267–Lys239 (DII–DII) maintained stable sb and hbss at 37 °C but lost over 20 % of both at 40 °C, and Lys321–Trp101 (DIII–DII; pc) disappeared at both elevated temperatures.

We then examined the isolated impact of pH variation at 37 °C (Figure 6C), identifying interactions absent at pH 7 but present at pH 5 or 6, along with significant frequency changes. In DENV–2, Gln256– Gly258 (hbsb; DII) was absent at pH 6 and pH 7, while Gln316–Phe108 (hbbb; DIII–DII) was absent at pH 5 and 7. Glu269–Lys241 (DII–DII; hbss and sb) lost over 10 % frequency at pH 6 and over 40 % at pH 7, while Glu314–Gly106 (hbbb; DIII–DII) occurred exclusively at pH 6, along with Glu314–Phe108 and Gly258–Ser255 (both hbbb). In DENV–3, unique pH 6 contacts included Asp98–Gly7 (DII–DI), Glu62– Lys200 (DII–DII), Glu247–Lys202, Glu247–Ser271, Glu255–Ser253, and Gly252–His259; Gly256–Ser253 was also present at pH 5. Most were within DII or involved multiple interaction types. Furthermore, Lys321–Trp101 (DIII–DII; pc) frequency dropped by 6 % at pH 6 and was absent at pH 7.

In summary, temperature and pH variations selectively perturb inter–domain interactions, promoting the loss of key contacts that weaken the dimer interface and favor monomerization. Reinforcing these interactions, both in solution and in the virion, represents a promising strategy for developing antibodies, small–molecule antivirals, and other therapeutic interventions to control dengue virus infection.

## 4 CONCLUSIONS

This study employed all–atom molecular dynamics simulations to show that pH and temperature act as synergistic physicochemical triggers that reshape the conformational landscape of the dengue–virus envelope protein (pE) in serotypes 2 and 3 (DENV–2 and DENV–3). Although the ectodomains are globally conserved, subtle differences in primary sequence–most notably in the spatial distribution of histidines–translate into serotype–specific dynamic responses and stability thresholds. DENV–2 displayed its greatest structural lability at pH 5 and 37 °C, conditions that mimic the human endosome, whereas DENV–3 became preferentially destabilized at pH 6 and 37 °C. Both serotypes also showed an intrinsic tendency to dissociate at neutral pH and 37 °C, underscoring their propensity for a monomeric state. Raising the temperature from 28 °C to 37 °C acted as a thermal “switch,” markedly expanding the conformational space and weakening the dimer interface through the loss or attenuation of salt bridges and hydrogen bonds.

A key finding was an asymmetric dissociation mechanism termed the “compensatory embrace.” As the protomers separate, recession of Domains I, II, or III in one monomer is counterbalanced by an advance of the corresponding domain in its partner, temporarily preserving intersubunit contacts and masking the initial separation. Essential motions reveal coupled rotation and rocking of these domains, identifying a tangible therapeutic target: disrupting this embrace should hamper the dimer–to–monomer transition and, consequently, viral fusion.

By correlating environmental cues with atom–scale rearrangements, the work refines current models of dengue–virus entry and delivers a detailed map of the interactions that underwrite dimer integrity. Selectively reinforcing these contacts—or directly inhibiting the compensatory–embrace mechanism—emerges as a rational route toward next–generation antivirals, broadly neutralizing antibodies, and vaccine candidates designed to stabilize pE and preclude membrane fusion.

## ACKNOWLEDGMENTS

This work was supported by Brazilian agencies: the National Council for Scientific and Technological Development-CNPq (grant 409272/2021-3), CAPES (88887.941707/2024-00), and the São Paulo Research Foundation (FAPESP) (grants: 2022/00347-0 and 2025/02641-0). Computational resources were provided by the National Laboratory for Scientific Computing (LNCC/MCTI, Brazil) for providing HPC resources for the SDumont supercomputer (Project antimicmd2) URL: http://sdumont.lncc.br, the Center for Scientific Computing (NCC/GridUNESP) of the São Paulo State University (UNESP), the “Centro Nacional de Processamento de Alto Desempenho em São Paulo (CENAPAD-SP), the Coaraci Supercomputer (Fapesp grant 2019/17874-0) and the Center for Computing in Engineering and Sciences at Unicamp (Fapesp grant 2013/08293-7), and the EuroHPC Joint Undertaking for awarding this project access to the EuroHPC supercomputer LEONARDO, hosted by CINECA (Italy) and the LEONARDO consortium through an EuroHPC Regular Access call. The authors acknowledge the use of ChatGPT (models o4 and o3) for improving the manuscript’s linguistic clarity and for assistance in plotting code development and troubleshooting. All outputs were rigorously reviewed by the authors, who assume responsibility for the final content.

## AUTHOR CONTRIBUTIONS

**Georcki Ropón-Palacios**: Conceptualization, Methodology, Software, Investigation, Formal analysis, Writing – Original draft preparation, Writing – Reviewing and Editing. **Ingrid B. S. Martins**: Writing – Original draft preparation, Writing – Reviewing and Editing. **Walter Rocchia**: Conceptualization, Writing – Original draft preparation, Writing – Reviewing and Editing. **Alexandre S. de Araujo**: Conceptualization, Project Administration, Writing – Original draft preparation, Writing – Reviewing and Editing.

## COMPETENTING INTERESTS

The authors declare no competing interests.

## DATA AVAILABILITY STATEMENT

Correspondence and materials requests should be addressed to the corresponding author Alexandre S. de Araujo.

